# Comparative morphology of the corpus callosum across the adult lifespan in chimpanzees (Pan troglodytes) and humans

**DOI:** 10.1101/2020.08.15.252205

**Authors:** René Westerhausen, Anders M. Fjell, Kristiina Kompus, Steven J. Schapiro, Chet Sherwood, Kristine B. Walhovd, William D. Hopkins

## Abstract

The human corpus callosum exhibits substantial atrophy in old age, which is stronger than what would be predicted from parallel changes in overall brain anatomy. To date, however, it has not been conclusively established whether this accentuated decline represents a common feature of brain aging across species, or whether it is a specific characteristic of the aging human brain. In the present cross-sectional study, we address this question by comparing age-related difference in corpus callosum morphology of chimpanzees and humans. For this purpose, we measured total midsagittal area and regional thickness of the corpus callosum from T1-weighted MRI data from 213 chimpanzees, aged between 9 and 54 years. The results were compared with data drawn from a large-scale human samples which was age-range matched using two strategies: (a) matching by chronological age (human sample size: n = 562), or (b) matching by accounting for differences in longevity and various maturational events between the species (i.e., adjusted human age range: 13.6 to 80.9 years; n = 664). Using generalized additive modelling to fit and compare aging trajectories, we found significant differences between the two species. The chimpanzee aging trajectory compared to the human trajectory was characterized by a slower increase from adolescence to middle adulthood, and by a lack of substantial decline from middle to old adulthood, which, however, was present in humans. Thus, the accentuated decline of the corpus callosum found in aging humans, is not an universal characteristic of the aging brain, and appears to be human-specific.

## Background

The corpus callosum is the major white-matter commissure (Schmahmann & Pandya, 2006) and has an important role in human perception and cognition (Gazzaniga, 2000). It supports the integration of complementary sensory information distributed across the two hemispheres (e.g., Genc, Bergmann, Singer, & Kohler, 2011; Steinmann et al., 2018; Westerhausen, Gruner, Specht, & Hugdahl, 2009), as well as the coordination of cognitive processing between the hemispheres (e.g., Chechlacz, Humphreys, Sotiropoulos, Kennard, & Cazzoli, 2015; Davis & Cabeza, 2015; Thiel et al., 2006). Furthermore, a series of neuroimaging studies suggest an association between variability in corpus callosum morphology and cognitive abilities (e.g., Danielsen et al., 2020; Dunst, Benedek, Koschutnig, Jauk, & Neubauer, 2014; Luders et al., 2007). Evidence for a common genetic origin for callosal size and intelligence measures (Hulshoff-Pol et al., 2006), as well as studies on callosotomy patients (Westerhausen & Karud, 2018), further underline the relevance of the corpus callosum for cognition.

However, in aging, the functional role of the corpus callosum is compromised by neurodegenerative processes affecting inter-hemispheric integration (van der Cruyssen, Gerrits, & Vingerhoets, 2020; Westerhausen, Bless, & Kompus, 2015). Histological studies have found a reduction in number and density of small myelinated axons (Hou & Pakkenberg, 2012) and degeneration of axonal myelin sheaths in old age (Bowley, Cabral, Rosene, & Peters, 2010; Peters & Sethares, 2002). These histological alterations are reflected in the findings of in-vivo diffusion MRI studies, which report a decrease in anisotropy and an increase in radial diffusion (i.e., orthogonal to the main fibre direction) in the aging corpus callosum (Hasan et al., 2009; Ota et al., 2006; Pietrasik, Cribben, Olsen, Huang, & Malykhin, 2020; Skumlien, Sederevicius, Fjell, Walhovd, & Westerhausen, 2018). Also, studies using high-gradient strength-based mapping of callosal fibre architecture suggest an aging-associated reduction in axon density (Fan et al., 2019). Morphometric assessment additionally suggests that not only the axonal composition of the corpus callosum is affected in aging, but also its midsagittal size. That is, atrophy both of midsagittal surface area (Doraiswamy et al., 1991; Hasan, Ewing-Cobbs, Kramer, Fletcher, & Narayana, 2008; Prendergast et al., 2015; Salat, Ward, Kaye, & Janowsky, 1997; Skumlien et al., 2018) and regional thickness (Danielsen et al., 2020) have been reported from middle to older adulthood. One often-overlooked feature of this old-age decline is that it is stronger than what would be predicted from the parallel decline in brain volume (Danielsen et al., 2020; Salat et al., 1997; Skumlien et al., 2018). That is, the ratio of corpus callosum to brain size declines in humans with advancing age rather than staying constant, suggesting over-proportionality of the decline, progressively disconnecting the two hemispheres. Individual differences in this decline appear to have consequences for cognitive functioning in older age (Danielsen et al., 2020; Salat et al., 1997), potentially by preventing compensatory recruitment mechanisms across hemispheres (Colcombe, Kramer, Erickson, & Scalf, 2005; Fling et al., 2011; Reuter-Lorenz & Cappell, 2008). Furthermore, clinical studies suggest that the relative decline is even more accentuated in Alzheimer patients than in controls (e.g., Frederiksen et al., 2011; Thomann, Wüstenberg, Pantel, Essig, & Schröder, 2006; Wiltshire, Foster, Kaye, Small, & Camicioli, 2005), additionally underlining the relevance of this over-proportional callosal decline for the understanding of human cognition in aging.

To achieve a better understanding of the aging corpus callosum, it is relevant to determine whether this over-proportional decline represents a common feature of brain aging across species, or whether it is a specific characteristic of human brain aging. To answer this question, here we compare age-related differences of corpus callosum morphology in humans and chimpanzees. Comparative studies with chimpanzees, as one of the closest living evolutionary and genetic relatives of humans, are especially important for this purpose, as they additionally offer the possibility to draw conclusions about human brain evolution (Rilling, 2014). Previous studies on brain aging suggest that chimpanzees show fewer signs of neurodegenerative processes in old age than humans (Chen et al., 2013; Herndon, Tigges, Anderson, Klumpp, & McClure, 1999; Sherwood et al., 2011). For example, while human aging is accompanied by an accentuated decline of brain white-matter volume from middle to old adulthood (e.g., Allen, Bruss, Brown, & Damasio, 2005; Raz, Ghisletta, Rodrigue, Kennedy, & Lindenberger, 2010; Walhovd et al., 2011), studies on chimpanzees reveal no, or comparatively milder, alterations of white-matter in old age (Chen et al., 2013; Sherwood et al., 2011). Thus, it is not surprising that the corpus callosum – as one of the major white-matter tracts – shows a similar dissociation between the two taxa. While studies on humans, as outlined above, find a prominent decline of callosal measures in old age, a preservation or even an increase in absolute or relative callosal area has been found in aging chimpanzees (Hopkins et al., 2016; Hopkins & Phillips, 2010). Thus, it may appear reasonable to conclude that over-proportional decline in hemispheric connectivity in aging is a human-specific phenomenon. However, a direct statistical comparison between chimpanzees and humans has not been conducted. Without such a comparison, it is difficult to ascertain whether human and chimpanzee trajectories are indeed different, or represent the same universal aging trajectory, which only affects humans more due to the longer lifespan, as previously suggested (Chen et al., 2013; Sherwood et al., 2011).

The objective of the present study was to compare the aging-related trajectory of corpus callosum morphology across the adult lifespan of chimpanzees to humans. For this purpose, the corpus callosum was analysed using MRI-based measures of total midsagittal area and thickness (relative to brain size), and general additive modelling (GAM) was utilised to fit and compare the aging trajectories. To account for substantial differences between the species in longevity and maturational events (e.g., onset of puberty, sexual maturity) across the lifespan (Robson & Wood, 2008), the direct comparison was done twice: (a) by comparing the chimpanzee with a human sample of the same chronological age range (chronological-age comparison), and (b) by using a human subsample with an age range that is adjusted to account for lifespan differences between the species (adjusted-age comparison). This approach allowed us to evaluate whether callosal aging trajectories are comparable, both considering absolute age and age relative to the expected lifespan.

## Method

### Chimpanzee Sample

This study sample consisted of 213 datasets from captive chimpanzees (*P. troglodytes*; 129 females, 84 males) covering an age range from 9 to 54 years (mean age ± standard deviation: 26.6 ± 10.3; for age distribution see Supplement Fig. S1). The data were retrieved from the National Chimpanzee Brain Resource (NCBR, www.chimpanzeebrain.org) and included all in-vivo MRI scans. Of note, of the original 227 datasets obtained, 14 datasets were excluded. For six, the MRI data quality was not sufficient to segment the corpus callosum, and for eight datasets, the quality of the segmentation was low or lesions in the corpus callosum were detected (for more details on the exclusion procedure see Supplement Section 2). Of the remaining study sample, 79 chimpanzees were housed at the Yerkes National Primate Research Center (YNPRC, Atlanta, Georgia) and 134 chimpanzees at the National Center for Chimpanzee Care (NCCC, Bastrop, Texas) at the time of MRI scanning.

The present data completely relied on available data collected prior to 2014 and no new data collection was conducted for the present study. The procedures of data collection were approved by the Institutional Animal Care and Use Committees at YNPRC and NCCC, and followed the guidelines of the Institute of Medicine on the use of chimpanzees in research. American Psychological Association guidelines for the ethical treatment of animals were adhered to during all aspects of this study.

### MRI scanning procedure and image acquisition

MRI scans followed standard procedures at the YNPRC and NCCC as described elsewhere (Hopkins et al., 2019) and were designed to minimize stress. In brief, the chimpanzees were first sedated (using ketamine (10 mg/kg) or telazol (3–5 mg/kg)) before being anesthetized with propofol (40–60 mg/(kg/h)). Then, the animals were transported to the MR imaging facility. After completion of the MRI acquisition, the animals were returned to their home facility and temporarily monitored in single housing, to ensure a safe recovery from the anesthesia, before returning to their social group.

T1-weighted images of 144 chimpanzees (6 from YNPRC, 138 from NCCC) were acquired with two 1.5T G.E. echo-speed Horizon LX MR scanner (GE Medical Systems), one at YNPRC and one at NCCC. Data was collected in transverse plane using a gradient echo protocol (repetition time, TR = 19.0 ms; echo time, TE = 8.5 ms; number of signals averaged = 8; scan matrix: of 256 × 256) with a reconstructed image resolution of 0.7 × 0.7 × 1.2 mm. The remaining 77 chimpanzees (all from YNPRC) were scanned on a 3.0-T Siemens Trio platform (Siemens Medical Solutions USA, Inc.). T1-weighted images were acquired using a 3D gradient echo sequence (pulse repetition, TR = 2300 ms; echo time, TE = 4.4 ms; number of signals averaged = 3; scan matrix of 320 × 320) yielding an 0.6 × 0.6 × 0.6 mm image resolution).

### Corpus callosum segmentation and measurements

Midsagittal callosal surface area and thickness were determined based on the T1-weighted images in native space performing the following processing steps. Firstly, to obtain a non-tilted midsagittal slice, individual images were coregistered to a template using rigid-body transformation (i.e., preserving size and shape of the corpus callosum) in SPM12 routines (Statistical Parametric Mapping, Wellcome Department of Cognitive Neurology, London, UK) and resampled to a 0.5 × 0.5 × 0.5 mm resolution. All resampled images were visually inspected to confirm a straight midsagittal plane as indicated by the longitudinal fissure forming a vertical line in coronal and axial views of the images. A midsagittal slice was identified using the criterion of minimal appearance of cerebral gray/white matter (lowest intensity) from regions adjacent to the longitudinal fissure. The cross-section of the corpus callosum was then manually traced on the midsagittal slice using MRIcron software (Rorden & Brett, 2000). Slides adjacent to the midline were used to inform the segmentation in cases where a delineation of callosal voxels from the fornix was required or where high intensity blood vessels were located close to the corpus callosum. For quality control, each tracing was rated on a three-point scale (0 = not usable, 1 = low, but acceptable quality, 2 = good quality; for details see Supplement Section 2). To account for data quality differences, statistical analyses including only the chimpanzee sample were done twice, once for all data of acceptable and good quality, and a second time for segmentations of good quality only. As can be seen in Supplement Section 12, the results of both analyses were comparable.

In a next step, the tip of the rostrum (defined as the inferior- or posterior-most voxel of the in-bend anterior callosal half) and the base of the splenium (ventral-most voxel in the posterior half) were identified on the callosal mask. Then, the mask was rotated so that an imagined line connecting rostrum tip and splenium base was orientated horizontally. The size of the callosal mask served as the measure of the midsagttal surface area for each individual. As reported previously, these manual segmentation steps yield interrater reliability estimates of *r*_ICC_=.86 and .96 (intra-class correlations calculated as two-way random effects, considering absolute agreement for a single measure) for midsagittal surface area (Danielsen et al., 2020).

To determine regional thickness, the outline of the callosal mask was automatically created and divided into a ventral and dorsal outline using the tip of the rostrum and the base of the splenium as dividing point (see also Westerhausen et al., 2016; Westerhausen et al., 2018). A midline between ventral and dorsal outline was determined as reference line for the thickness measurements. That is, 100 support points spaced equidistantly on the two outlines were created and the midline coordinates were calculated as average coordinates of the two corresponding support points. The resulting midline was resampled into 60 equidistant points that marked the location of the thickness measurement. Callosal thickness was defined as the distance between the ventral and dorsal outline orthogonal to the midline at these points. The number of 60 measurement points was chosen, as it provides a sufficiently high density of sampling points to capture the structure of the corpus callosum, while not excessively inflating the number of statistical tests (Danielsen et al., 2020; Westerhausen et al., 2016).

As the aim of the present analysis was to examine the proportionality of corpus callosum age-related differences, both area and thickness measures were divided by forebrain volume (FBV, see next section). However, as suggested by Smith (2005), we converted FBV before the division so that it had the same unit as the respective callosal measure, as only under this condition the ratio is expected to be constant if corpus callosum and FBV are proportional to each other. In other words, FBV was converted to a unit that stays proportional to callosal area and thickness, respectively, if the brain with changing size maintains geometric similarity. In practice, we raised FBV to the power of 2/3 (i.e, FBV^0.666^) to calculate the ratio with area, and to the power of 1/3 (i.e, FBV^0.333^) to calculate the ratio with thickness (for a detailed explanation refer to Smith, 2005). The resulting ratios are hereafter referred to as relative callosal area and relative callosal thickness. Constant relative area and thickness across the lifespan would indicate proportionality of the developmental differences in the corpus callosum, while any positive or negative deviation reflects over-proportional increase or decline, respectively.

### Brain-size extraction

FBV was selected to account for brain size differences. FBV was preferred over measures of total intracranial volume, as the corpus callosum is formed from axons originating from the two cerebral hemispheres (Schmahmann & Pandya, 2006), and brain structures irrelevant for the corpus callosum (e.g., brain stem, cerebellum) are excluded. For this purpose, a custom mask was created covering the supra-tentorial brain in standard space defined by the chimpanzee template. FBV was then determined for each data set in three steps. Firstly, using SPM12 brain segmentation routines, gray- and white-matter maps were created in native space (using the chimpanzee template tissue probability maps from (Vickery et al., 2020)). Then, the standard FBV mask was transferred to the individual brain by using the same transformation parameters used when creating the tissue segmentations in native space. Finally, FBV was determined as the sum of gray- and white-matter probabilities within the mask in native space. Thus, the resulting FBV estimate did not include CSF compartments.

### Human comparison data

For comparison of the chimpanzee with human corpora callosa, we included data from a large, mixed longitudinal and cross-sectional sample including 1867 datasets from 1014 (608 female) healthy participants (Danielsen et al., 2020). The age range of the sample spanned from 4 to 93 years (mean: 33.8 ± 24.4 years). The corpus callosum measurements in this sample were extracted using a mostly identical approach to the extraction of the chimpanzee data as described above (for details see (Danielsen et al., 2020), and Supplement Section 3). The only difference was that the segmentation of the corpus callosum was based on an automated initial white-matter segmentation step (obtained using standard SPM12 routines), rather than the manual approach chosen here. The difference in approach was necessary as the chimpanzee data frequently showed hyper-intensity artifacts of arteries on the midline, which prevented a reliable initial segmentation based on the white-matter maps. However, also for the human sample, the initial segmentation was followed by manual adjustment, so that the final callosal mask was carefully created and adjusted by the examiners. Thus, we regard the results of the two approaches as equivalent.

The lifespan and developmental milestones of humans and chimpanzees differs substantially (Robson & Wood, 2008), so that the age range of a human sample included in the comparison has to be considered carefully. We here chose to draw two comparison subsamples from the above-described human sample. That is, (a) a same chronological age subsample using the identical age range in humans and chimpanzees (i.e., 9 to 54 years), and (b) an adjusted-age subsample in which chimpanzee age is transferred into a human-age equivalent. Concerning the latter, we compared the timing of certain life events in the two species to determine a factor to transfer chimpanzee age into a human age equivalent. Comparing the onset of puberty (Behringer, Deschner, Deimel, Stevens, & Hohmann, 2014; Kelsey et al., 2014), sexual maturity (Robson & Wood, 2008), as well as the maximal lifespan (Hill et al., 2001; Robson & Wood, 2008) as reported in the literature (see Supplement Section 4 for details), we concluded that a factor of 1.5 offers a reasonable approximation for this transformation. That is, the here studied age range of chimpanzees between 9 and 54 years was considered comparable to a human age range of 13.5 to 81 years. As the chimpanzee data were cross-sectional, both human comparison subsamples were additionally restricted to the first dataset acquired in the respective age range.

The resulting chronological-age comparison subsample included 562 participants (364 females). The participants had a mean age of 27.8 ± 12.1 years and the exact age range was 9.0 to 53.8 years. The MRI of 313 and 249 of these datasets had been acquired on a 1.5 Tesla Siemens Avanto and on a 3 Tesla Siemens Skyra system, respectively. The adjusted-age subsample consisted of 664 participants (430 females) with a mean age of 37.2 ± 17. 6 years (exact range: 13.6 to 80.9 years) and included 394 Avanto and 270 Skyra scans.

### Statistical analyses

The lifespan trajectories of relative area and thickness in chimpanzees were fitted with generalized additive models (GAM) using the “mgcv” package (v1.8-31; (Wood, 2017) using R 3.6.2). That is, Age was smoothed using cubic regression splines with 6 knots as a basis dimension. The participants’ Sex and Scanner Type (Siemens 3T vs. GE 1.5T) were added as covariates. To test for sex differences of the developmental trajectories, these analyses were followed by a second analysis step including terms for the modulation of the Age trajectories by Sex (i.e., set up to test for a deviation of the male from the female trajectory). Model estimation was done using the restricted maximum likelihood (REML) method. Example R code calling the gam fitting function can be found in Supplement Section 5.

The direct comparison of the lifespan trajectories between the taxa was calculated separately for the comparison of chimpanzees with the chronological-age and the adjusted-age human subsamples. Of note, for the comparison with the adjusted-age group, the age of the chimpanzee sample was multiplied by the factor 1.5 (as indicated above) to facilitate the direct comparison. In both cases, the model was set up to test for the deviation of the chimpanzee from the human trajectory by including the Age by Group term in addition to the Age term. The model included Sex and Scanner Type (4 levels, reflecting the scanner types used in the chimpanzee and human samples) as additional predictors. A Group predictor was not included, as it would be collinear to the Scanner Type predictor.

Across all analyses, the effect size was expressed as proportion of explained variance (ω^2^). Considering the segment-wise thickness analyses, the p-values were adjusted for multiple comparison to yield a false discovery rate (FDR) of 5 %, and an extent threshold of 3 segments was additionally applied to remove spurious effects. Where relevant for the discussion, we also determined the end of growth and the beginning of decline of the fitted trajectories, defined as the last estimate above and the first estimate below zero, respectively. For this purpose, the derivatives (i.e., slope) of the fitted trajectories were determined, and the confidence bands (95%) around the derivative trajectory were used to decide where the slope deviated from zero.

### Data-availability statement

All tabulated data as well as R scripts for reproduction of the present results are available on an Open Science Framework platform (https://osf.io/5yzb3/). The raw MRI data of the chimpanzee sample can be requested from the National Chimpanzee Brain Resource (www.chimpanzeebrain.org). Requests for accessing the raw human MRI data need to be directed to the principal investigators of the original LCBC studies (contact details can be found here: www.oslobrains.no). Here, data sharing requires appropriate ethical and data-protection approvals, within a collaborative project.

## Results

### Descriptive statistics of corpus callosum measures

The averaged outline of the chimpanzee corpus callosum and the thickness profile are shown in Fig. 1A and 1B. The mean absolute midsagittal callosal area in the chimpanzees was 263.3 ± 41.8 mm^2^ and ranged from 145.0 to 374.4 mm^2^ (see histogram in Fig 1C). The mean relative callosal area was 0.063 ± 0.009 (range: 0.032 to 0.089).

**Fig. 1.**
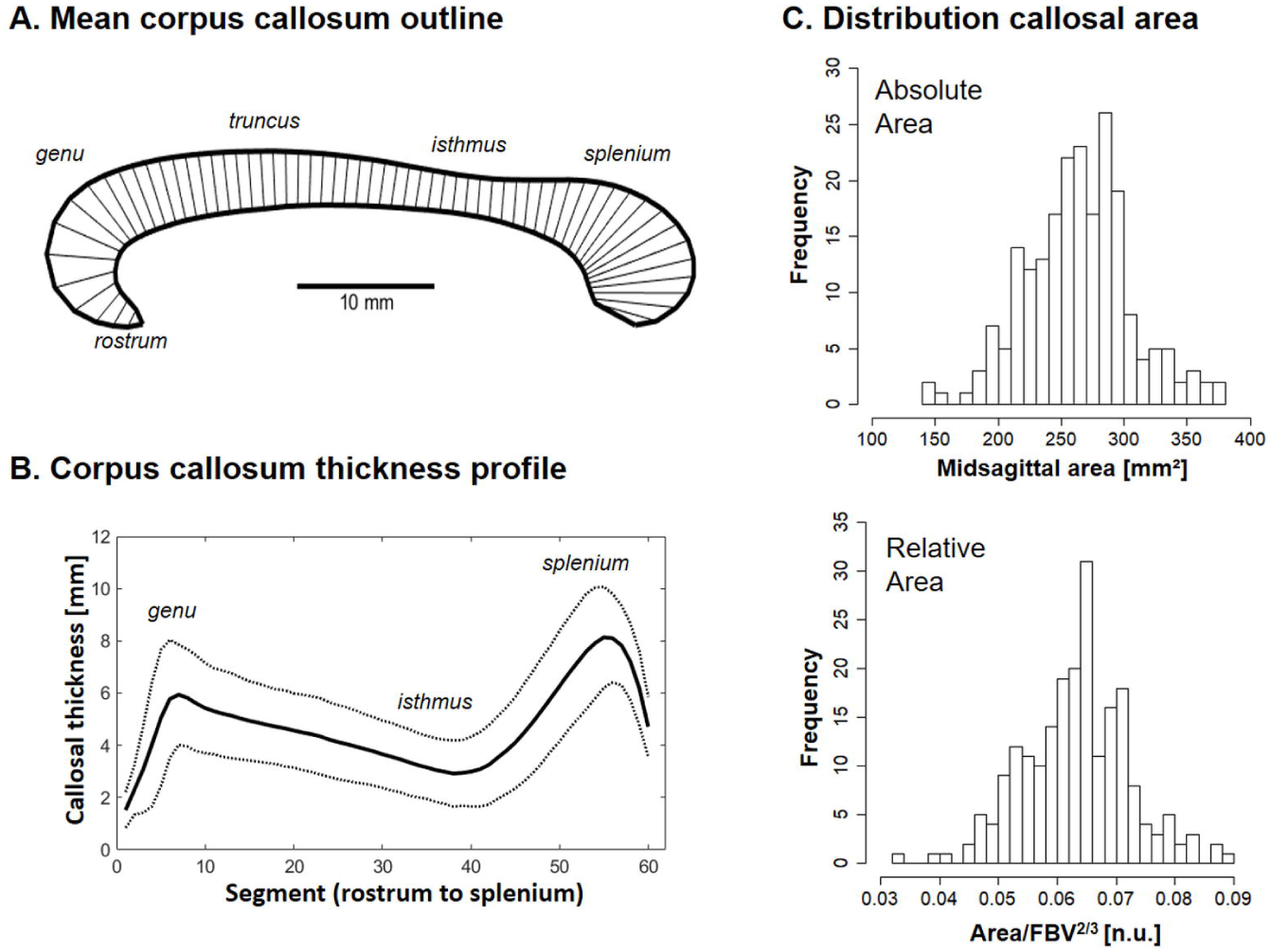
The chimpanzee corpus callosum. Panel A shows the mean outline of the corpus callosum averaged across all 213 individuals of the study sample. The lines within the corpus callosum represent the 60 thickness measurement points (segments). Panel B presents the mean (± 95% confidence limits) thickness for each of the segments. Panel C shows the histograms for mean absolute (top) and relative area (bottom) of the corpus callosum.

### Trajectories of midsagittal callosal measures in chimpanzees

The age trajectory of relative callosal area is depicted in Fig. 2. The fitted age trajectory was significant with edf (effective degrees of freedom) of 2.06 (*F* = 8.33, p <.0001), explaining 6.6% of the variance in the data. The slope of the trajectory was positive in young adults, and decreased until the age of 30.8 years, where it no longer deviated from zero, marking the end of callosal growth. No decline in corpus callosum area was observed, as the slope did not deviate negatively from zero in the studied age range .The main effect of Sex was not significant (*t*(207.9) = −1.33, *p* = 0.18, ω^2^ < .001). The follow-up analysis also did not find a deviation of the male from the female age trajectories (edf = 1.00, *F* < 1, *p* = 0.46, ω^2^ < .001, see Supplement Fig. S2). An analysis of absolute area yielded comparable trajectories (see Supplement Section 7).

**Fig 2.**
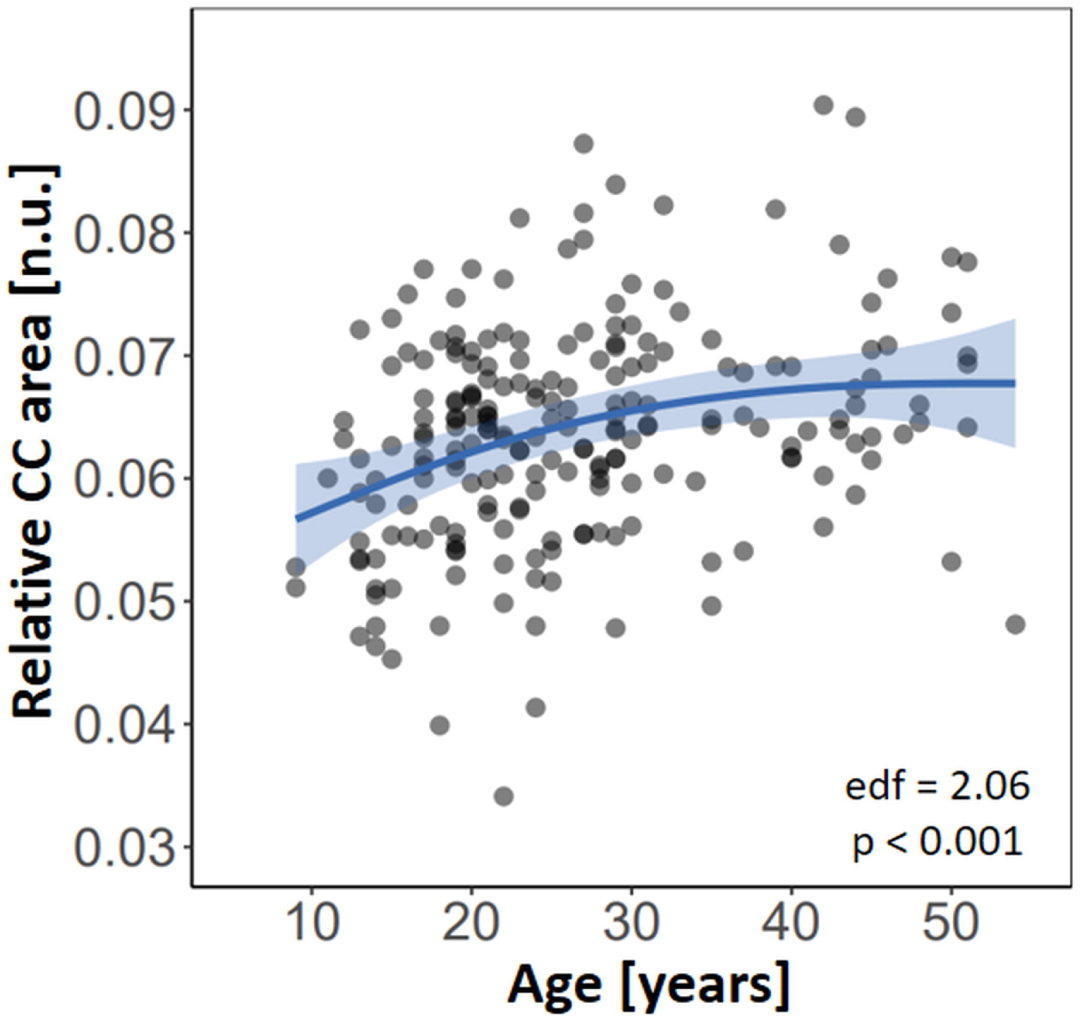
Relative midsagittal area of the chimpanzee corpus callosum across the adult lifespan. The blue line represents the GAM fit (shading: 95% confidence band) of the prediction of relative area by Age (using Scanner type and Sex as covariate). Note: the ratio (y-axis) does not have a unit (n.u.), as the units of numerator and denominator cancel out.

As shown in Fig. 3, a cluster of significant age trajectories (at FDR = 0.05) was found in the genu of the corpus callosum. The age effect was significant in segments 11 to 15 (edf: between 2.22 and 2.50) explaining 4.3 to 6.8% of the variance. The slope of the mean cluster trajectory was significantly above zero until the age of 29 years, marking the end of growth, and no significant decline was observed thereafter. No significant deviation in the trajectories of males compared to females was detected (for all segments *p*_*FDR*_ >.43). However, a significant main effect of sex was found in segments 33 to 35 (truncus subregion), indicating larger relative thickness in female compared to male chimpanzees (all *p*_*FDR*_ <0.048; all *t* < – 2.91, all ω^2^ > .034). A topography of the edf of the age trajectory, irrespective of significance, can be found in Supplement Fig S4. Segment-wise statistics can be found in Supplement Tables S1.

**Fig 3.**
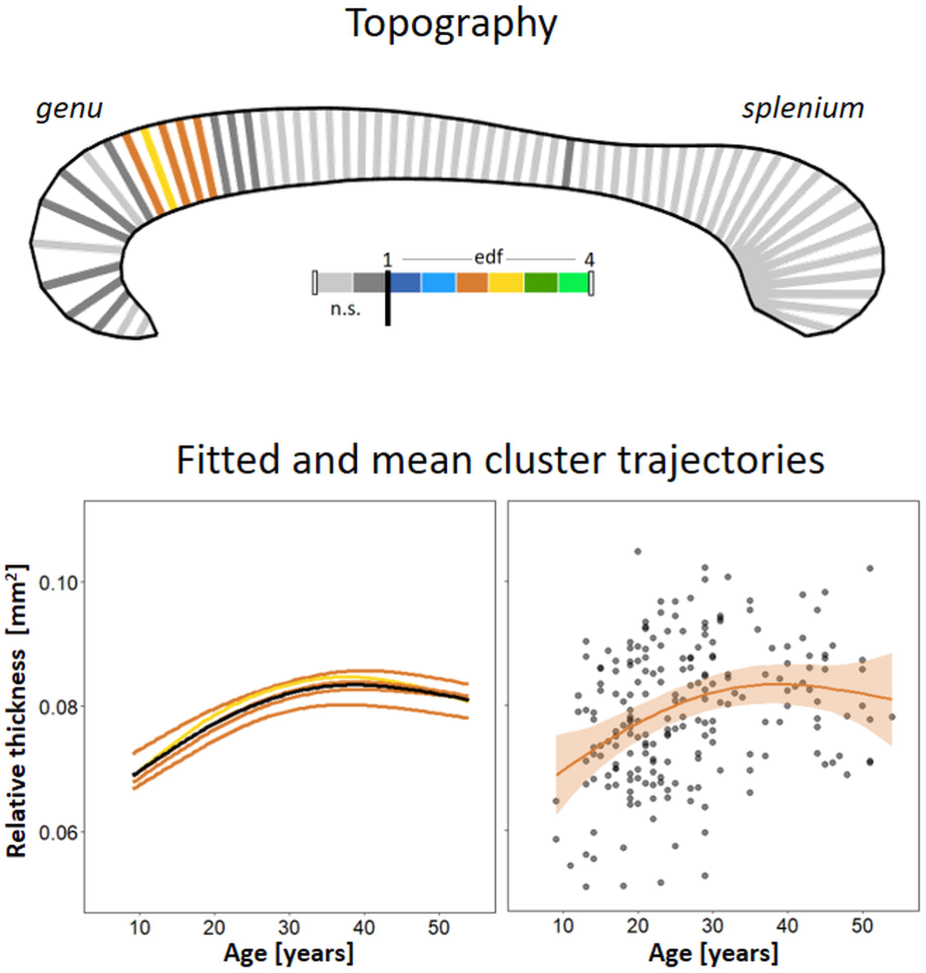
Relative thickness of the chimpanzee across the adult lifespan. Upper panel presents the topography of the age effect. The colouring codes the edf of the fitted trajectory in the segment. Segments depicted in grey do not show a significant effects after FDR correction. Dark grey segments show significance before FDR correction. The lower panels shows the fitted trajectories separately for all significant segments (left) and the mean trajectory of all significant segments, its 95%-confidence band, and the individual data points (right). Note: the ratio (y-axis) does not have a unit (n.u.) as the units of numerator and denominator cancel out.

### Comparison of chimpanzee and human trajectories

Comparing the chimpanzee sample with the human subsample of same chronological age revealed a significant deviation of the trajectories in which relative corpus callosum area develops (see Fig 4). That is, while the age effect in the human sample with a complexity of edf = 3.10 (*F* = 7.85, p<.0001, ω ^2^ =.03) displays an inverted-u-shaped trajectory, the chimpanzees deviated from this trajectory with a more or less linear (positive) trajectory (edf = 1.29, *F* = 4.71, p = .03). As shown in Fig. 4, the deviation was driven by a negative deviation in the younger age subjects, indicating that the chimpanzee trajectory runs below the human trajectory (up to approx. 20 years). In older age subjects, the deviation is positive, but only around the age of 40 years does the 95% confidence band not include zero. However, the deviation trajectory explained under 1% of the variance in the data (ω^2^ =.006).

**Fig. 4.**
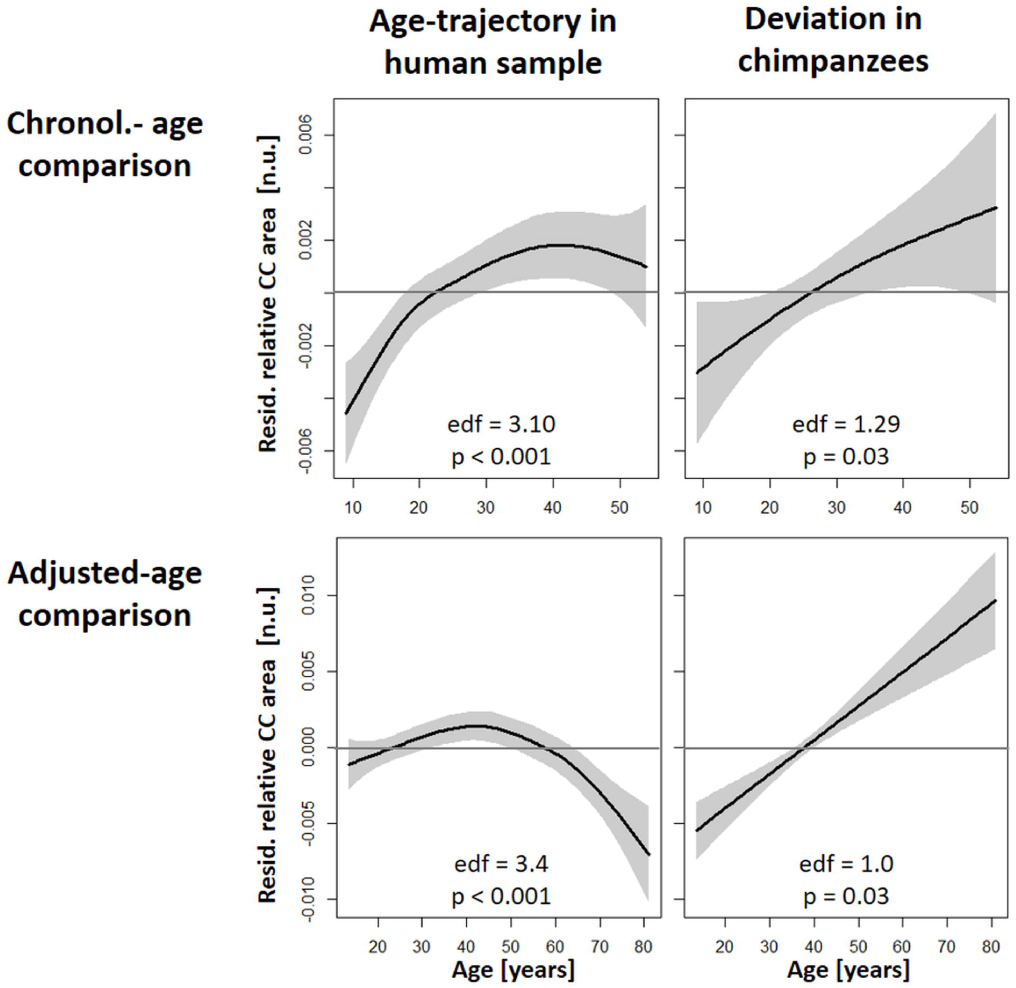
Direct comparison of chimpanzee and human age trajectories of relative callosal area. Graphs in the upper row presents the direct comparison of the chimpanzee trajectory with the human trajectory of the chronological-age reference group. That is, the left graph shows the fitted trajectory in humans, while the right trajectory depicts the deviation of the chimpanzee trajectory. The graphs in the lower row present the results for the comparison with the adjusted-age human reference group. Here, the age range in chimpanzees was transformed to a human age-equivalent (for details refer to method section).

Comparing the chimpanzees with the age-adjusted subsample further highlighted the above effects. While the human trajectory was fitted with an edf = 3.4 (*F* = 6.62, *p* <.0001, ω^2^ =.02) describing an inverted u-shaped trajectory, the deviation in chimpanzees was linear (*F* = 36.7; edf = 1.0, *p* <.0001), and now explained 4% of the variance in the data. As can be seen in Fig. 4, at younger ages, the chimpanzees deviate negatively, and at older ages positively, from the human trajectory, indicating the less steep increase and slower/no decline, respectively, in the chimpanzee sample.

Figure 5 shows the results of the chronological-age and adjusted-age comparisons of the segment-wise relative thickness analysis. The chronological-age comparison revealed three clusters in which the deviation trajectory was significant. The clusters were located in the rostrum (segments 01 to 05), genu (segments 11 to 18) and isthmus (segments 40 to 46). Although the deviation trajectories differ in their complexity between the clusters (i.e., edfs were higher in the rostrum), the three clusters had in common that at young ages (up to early 20s), the deviation from the human trajectory was negative, while at older ages, it was positive (see graphs in Fig. 5A). The adjusted-age comparison indicated significant deviation trajectories in almost all segments, except for segments of the ventral splenium (see Fig. 5B). Throughout dorsal genu, truncus, isthmus, and splenium, the deviation was linear (edfs between 1 and 1.5), but in the rostrum and ventral genu, higher edfs were found (up to 3.38; cf. Supplement Table S2). However, the general pattern indicated a negative deviation at young ages (up to an adjusted age of ca. 30 to 40 years), and a positive deviation in older age in chimpanzees, compared to the human reference trajectory. Compared with the chronological-age analysis, the deviation trajectories were not only more widespread, but the variance explained was also higher in the age-adjusted analysis, reaching up to 5-6% in the genu (cf, Supplement Table S2). This accentuation was driven by the human sample, which now included older participants, for which a significant reduction in callosal thickness can be observed throughout all corpus callosum subregions (see fitted trajectories, Fig. 5B, and edf topography in Supplement Fig. S5).

**Fig 5.**
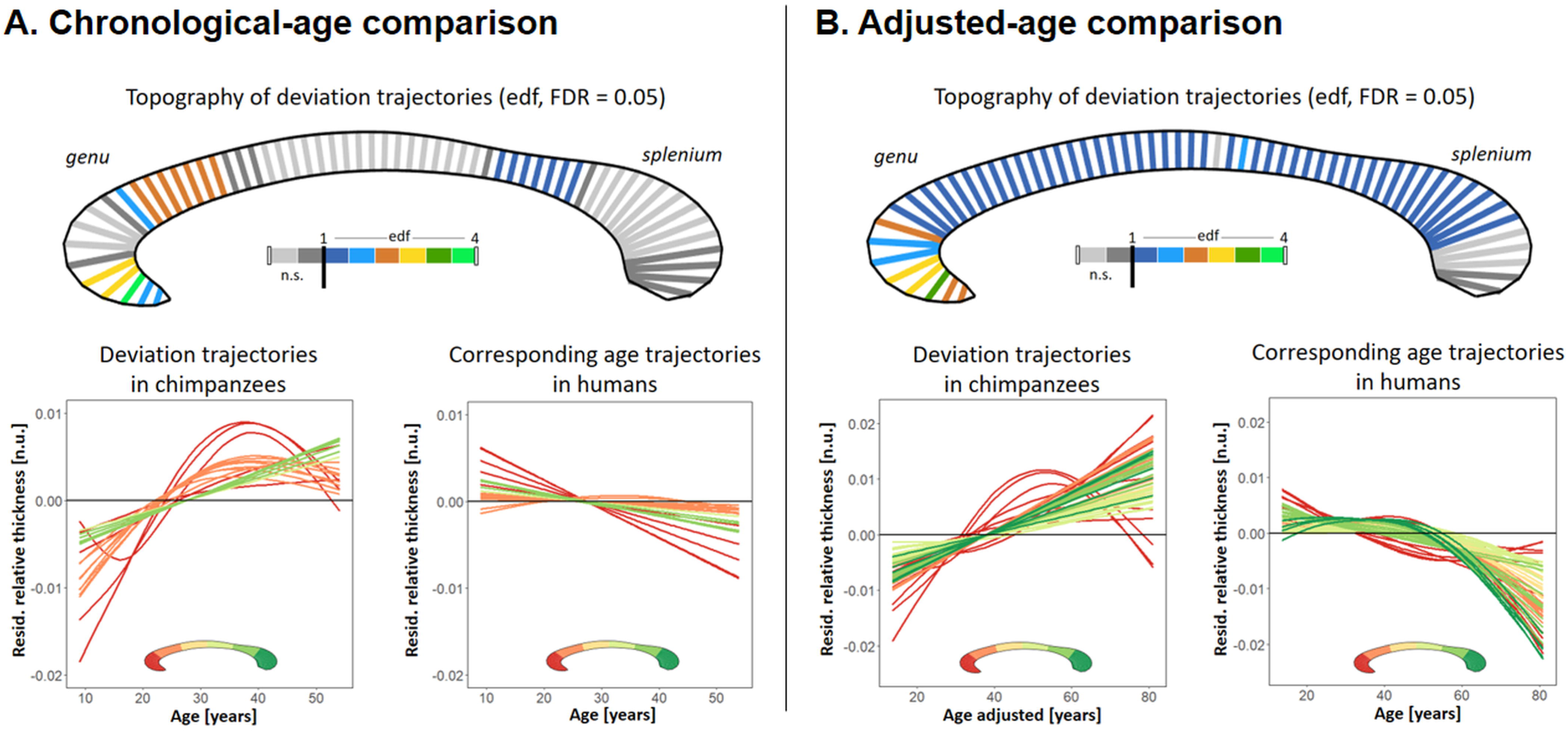
Direct comparison of chimpanzee and human age trajectories of relative callosal thickness. Panel A presents the comparison with the chronological-age human reference sample; panel B the comparison with the adjusted-age sample. In both panels, the upper row illustrates the topography of the deviation trajectory (colour-coded according to its edf) of chimpanzees compared to the human trajectories (grey segments: deviation not significant after FDR correction to 5%). Note, the callosal outline represents the mean outline of human and chimpanzees. The graphs in the lower row show the chimpanzee’s deviation trajectories for significant segments and the corresponding age trajectories of the human reference sample (right). Here, the colour codes the spatial location of the segment in accordance with the corpus callosum colour map insert at the bottom of each graph. The topography of the edfs of the human aging trajectory can be found in Supplement Figure S5.

## Discussion

Comparative neuroimaging studies of humans and chimpanzees have improved our understanding of brain evolution by identifying human-specific brain characteristics (Rilling, 2014). Recent morphometric studies suggest that one such characteristic is that the trajectories of brain ageing differ between the two species. The human brain exhibits a continuous reduction of cortical gray matter across the lifespan from late childhood to old age (Allen et al., 2005; Walhovd et al., 2016; Walhovd et al., 2011), and an accentuated decline of white matter from middle to old adulthood (Allen et al., 2005; Raz et al., 2010; Walhovd et al., 2011). Studies of the chimpanzee brain reveal no, or only comparatively mild, volumetric changes in total brain, gray, or white matter with advancing age (Chen et al., 2013; Herndon et al., 1999; Leigh, 2004; Sherwood et al., 2011; Vickery et al., 2020). Expectedly, similar dissociation between the two species are also found regarding the corpus callosum. While studies on humans report a prominent decline of midsagittal callosal area (Doraiswamy et al., 1991; Hasan et al., 2008; Prendergast et al., 2015; Raz et al., 2010; Skumlien et al., 2018) or thickness (Danielsen et al., 2020), preservation of callosal area has been reported in aging chimpanzees (Hopkins et al., 2016; Hopkins & Phillips, 2010). The present study, while confirming the general differences in aging trajectories of the corpus callosum of the two species, offers a series of important clarifications and novel data to the literature.

Firstly, the present findings specify the shape of the aging trajectory in chimpanzees and confirm the differences between the species in a direct comparison. That is, the utilized GAM fitting suggests an asymptotic growth curve of (relative) corpus callosum area in chimpanzees. The increase that is observed in younger age decelerates with aging, reaching a constant level above the age of 30 years, while showing no significant changes in older age (see Fig. 1). This contrasts with previous analyses, which report a linear trajectory of development, suggesting an ongoing increase in relative area into old age (Hopkins et al., 2016; Hopkins & Phillips, 2010). This difference in the reported trajectories may be attributed to the fact that previously used linear or cubic polynomial fitting approaches are rigid, as the entire age range determines the results (Fjell et al., 2010), while in this study, we used GAMs, by means of smooth terms, which optimise curve fitting to better fit “local” data points (Sørensen et al., 2020). The preservation of callosal connectivity in aging chimpanzees not only represents an apparent deviation from the human trajectory, the deviation is also statistically significant in direct comparison. For example, considering the age-adjusted comparison, the age trajectory of the human sample shows the clear atrophy of relative area (Fig. 4) expected from the literature (e.g., Salat et al., 1997; Skumlien et al., 2018). The chimpanzee data deviate linearly from this decline above the age of ca. 40 years (adjusted age), implying an increasing gap between the two trajectories with advancing age, and reflecting the lack of a decline in chimpanzees as indicated above.

Secondly, using callosal thickness measures, regional variation in the age trajectories and their deviation from human trajectories were found. Significant age trajectories were localised selectively in a cluster of segments in the anterior third (or genu) of the corpus callosum, while for all other segments, no significant associations were detected. The shape of the trajectory within this cluster was comparable to the trajectory of the relative area: an increase in thickness in young age reaches its plateau around the age of 30 years. Axons located in this section of the corpus callosum are thought to connect prefrontal, premotor, and cingulate cortices of both hemispheres, as converging evidence across primate species suggests (i.e, post-mortem tracing studies in rhesus monkeys, (Schmahmann & Pandya, 2006), or chimpanzee (Phillips & Hopkins, 2012) and human diffusion-imaging tractography (Archer, Coombes, McFarland, DeKosky, & Vaillancourt, 2019; Chao et al., 2009)). Thus, it is tempting to speculate that the observed accentuated age effects in the genu of chimpanzees might be related to a) ongoing development in these connected cortical regions up to middle adulthood, and b) higher order motor planning and cognition. The lack of a substantial age effect in the remaining segments indicates that relative callosal thickness is statistically stable within this age range, which may suggest that the major ontogenetic growth of callosal thickness takes place below the lowest age range covered in the present study (see Sakai et al., 2017). The adjusted-age comparison with human trajectories, yields significant (mostly linear) deviation of the chimpanzee from the human trajectories across most segments of corpus callosum. In humans, the relative thickness measures of most segments are stable until about an age of 40 to 50 years, followed by a strong thickness reduction in older age (see Fig. 5). Chimpanzees deviate positively from this trajectory in older age, indicating that a comparable old-age thickness reduction is missing. Segment trajectories of the ventral genu and rostrum seem to differ from this general pattern, as here chimpanzees deviate in trajectories that are more complex. However, this effect is mainly driven by the human sample showing a constant decline across the included age range in the anterior corpus callosum (Danielsen et al., 2020).

Thirdly, the above findings complement previous comparative imaging studies that report an enhanced aging-related decline of white matter in humans as compared to chimpanzees (Chen et al., 2013; Sherwood et al., 2011). Sherwood et al. (2011) did not find a significant reduction in total or frontal white-matter volume in chimpanzees in older age, while a clear age-related decline of both measures was reported for humans, but only beyond the lifespan of chimpanzees. This led the authors to conclude that aging trajectories are similar in the two species, but only humans would get sufficiently old to show significant white-matter deterioration. Chen et al. (2013) specified these findings by showing decline of whole white-matter anisotropy in both chimpanzees and humans. The decline was observed from the same absolute age (ca. 30 years) and followed comparable aging trajectories. From this, the authors concluded that the aging trajectories of both species are in general comparable considering absolute age, but the longer lifespan would allow human white matter to decline longer. The present study, in line with Sherwood et al. (2011), does not find a significant decline in callosal white matter in chimpanzees, presumably, as the planimetric measures used are less sensitive to microstructural change revealed by Chen et al. (2013) using diffusion imaging. However, the direct comparison of humans and chimpanzees of same chronological age, revealed differences in the age trajectories of both total area and regional thickness (see Figs. 4 and 5). Although these differences were present across the studied age range, the small, but significant, positive deviation of the chimpanzee from the human trajectories from middle to old adulthood contrast the findings of comparable aging trajectories reported by the above discussed earlier studies (Chen et al., 2013; Sherwood et al., 2011). Thus, while agreeing with both studies that differences in longevity certainly allow white matter of the human brain to decline longer, we provide evidence that the trajectories of the corpus callosum decline in old age actually deviate significantly before lifespan differences manifest themselves.

Fourthly, while previous comparative studies focussed on effects in older age, the present study indicates that species differences in the developmental trajectories also exist from young to middle adulthood. That is, relative total area and genu thickness measures indicate an end of callosal growth around the age of 30 years in chimpanzees. In humans, “peak” corpus callosum is reached earlier, both when considering the actual age and, certainly, when transforming the chimpanzees’ estimate into human age-equivalent (i.e., ca. 45 years). For comparison, in the present human sample, an end of growth for relative area was estimated to be at ca. 23 years (see Supplement Section 13), which is well in accordance with previous studies suggesting maximum area being reached in the early twenties (Pujol, Vendrell, Junque, Marti-Vilalta, & Capdevila, 1993). Likewise, thickness measures of the genu segments here shown an age effect in chimpanzees, and reach maximum values between 17 and 24 years (Danielsen et al., 2020). This observation is also confirmed by the direct comparison, as both adjusted-age and chronological-age comparisons show a negative deviation of the chimpanzee from the human trajectories in the younger age range (see Fig. 5). In fact, previous studies suggest that these species differences in callosal growth rate can also be found below the age ranges in this study. During infancy, chimpanzees show roughly a doubling of total callosal area within the first 24 month (Sakai et al., 2017), while human corpora callosa appear to quadruple in the same period (Rakic & Yakovlev, 1968; Tanaka-Arakawa et al., 2015; Vannucci, Barron, & Vannucci, 2017). Thus, growth of the chimpanzee corpus callosum seems to be slower and protracted compared to the human development. One might speculate, that the accentuated growth is related to interspecies difference in relative brain size at birth. While chimpanzee’s brain at birth has approximately 36% of the adult size, in humans it is 25% (Robson & Wood, 2008), thus requiring a stronger postnatal growth to reach adult brain size. Nevertheless, this does not account for the relative longer growth period of the corpus callosum in chimpanzees.

Taken together, human and chimpanzee trajectories of callosal development differ across the lifespan. Compared with the trajectories found in chimpanzees, the human trajectory appears accentuated: setting off in a steeper angle during infancy and young adulthood, reaching the peak earlier, and showing significant decline in old age. Studies in human and other primate species suggest that the growth of the corpus callosum during infancy and childhood likely reflects an increase in axon diameter and myelination, but less likely an increase in the number of axons (Clarke, Kraftsik, Van der Loos, & Innocenti, 1989), as the number of axons is subject to extensive postnatal pruning (Innocenti & Price, 2005; LaMantia & Rakic, 1990). Aging-related callosal atrophy is likely driven by the loss of small myelinated axons (Bowley et al., 2010; Hou & Pakkenberg, 2012; Tang, Nyengaard, Pakkenberg, & Gundersen, 1997) but also alterations of myelin sheaths may contribute to the aging process (Bowley et al., 2010; Peters & Sethares, 2002). However, little is known about differences in the fibre composition of the corpus callosum between primate species (Caminiti, Ghaziri, Galuske, Hof, & Innocenti, 2009; Hopkins et al., 2012; Phillips et al., 2015), let alone about differential age-related changes in callosal axons between the species. Thus, without future comparative studies, it remains impossible to determine which neuronal mechanism underlies the observed macrostructural developmental differences reported here. Assuming a lifespan perspective, one might speculate that the mechanism underlying the relatively pronounced growth of the human corpus callosum from infancy into adulthood also might render the corpus callosum more vulnerable for deteriorations approaching older adulthood.

Finally, sex did not significantly modulate the age trajectories found in area and thickness analysis of the chimpanzee corpus callosum. This observation is in line with the findings in the present human sample for which previous analyses also did not show any substantial sex differences in callosal thickness trajectories (Danielsen et al., 2020). Sex differences in the mean relative area (or absolute area, see Supplement Section 7) were also not significant in our chimpanzee sample, thereby confirming the results of a previous analysis of a largely overlapping sample (Dunham & Hopkins, 2006; Hopkins et al., 2016). In order to evaluate what effect sizes can be excluded considering the present sample size, we supplemented the present analyses with a sensitivity power analysis. The results suggest that population sex effects of d > 0.35 (ca. 3% explained variance) may be excluded with acceptable test power (.80, at α = 0.05, one-tailed, using G Power software, (Faul, Erdfelder, Buchner, & Lang, 2009)). Regarding relative callosal thickness, however, we found larger relative thickness in female compared to male chimpanzees, in three segments located in the posterior truncus section. The region likely connects pre-motor and motor cortices (Phillips & Hopkins, 2012; Schmahmann & Pandya, 2006) so that the association we found might be related to sex differences in manual performance previously found in chimpanzees (Hopkins, Russell, Schaeffer, Gardner, & Schapiro, 2009). Overall, however, sex differences in the chimpanzee corpus callosum appear to be small and regionally restricted, which is arguably comparable to the typical findings in humans. That is, while human studies suggest that the female corpus callosum area relative to brain size is larger than the male, the pooled effect size reported in a meta-analysis is small, explaining ca. 1% of the variance in the data (Smith, 2005). Likewise, regarding the thickness measure recent studies fail to report substantial sex differences in human samples (Danielsen et al., 2020; Luders, Narr, Zaidel, Thompson, & Toga, 2006; Luders, Toga, & Thompson, 2014).

The present study comes with a set of limitations, which deserve discussion. Firstly, the chimpanzee sample (and consequently the human comparison sample) consisted of cross-sectional data, which cannot be utilized to conclusively determine age-associated changes in brain structure (Pfefferbaum & Sullivan, 2015; Salthouse, 2011). That is, the trajectories we find represent differences between individuals of various ages and not changes within individuals, so that, for example, potential cohort effects cannot be distinguished from “true” age-related change on an individual level. Comparing the present cross-sectional analysis of the human callosal thickness data with the previous mixed-effect analysis also including longitudinal measurements suggest that the results are comparable (cf. Danielsen et al., 2020). However, there is a major cohort difference between older and young chimpanzees: the vast majority of the older chimpanzees were born in the wild, while chimpanzees under 40 were usually born in captivity. Thus, here found age-related differences are potentially confounded with differences in early-life experiences, which cannot be statistically controlled given the present data. Secondly, we here utilized macro-anatomical measures of the corpus callosum and consequently were not likely to capture age- or species-related differences in the mircrostrucural composition of the corpus callosum. As shown by Chen et al. (2013), diffusion imaging might be able to detect age-related deterioration of white matter earlier than volumetric measures. Histological analyses suggest, however, that midsagittal area is a valid predictor of the number of myelinated axons in the corpus callosum (Aboitiz, Scheibel, Fisher, & Zaidel, 1992; Hou & Pakkenberg, 2012; Riise & Pakkenberg, 2011) so that we believe that our results can be interpreted to reflect differences in the strength of the callosal connectivity across the lifespan within and between the species. Arguably, the present findings would be strengthened, if future studies found similar aging trajectories of corpus callosum development in histological or diffusion studies. Thirdly, chimpanzees and humans participants were not scanned on the same MRI scanner platforms and scanning sequences, so that a direct comparison of mean callosal measures is confounded by difference in image acquisition. We consequently refrained from discussing mean differences in the callosal measures between the species. However, we have no reason to believe that the comparison of the aging trajectories across species is affected by the scanner differences, as the shape of the trajectory is fitted within group and we do not expect any age by scanner interaction.

## Conclusion

The present findings indicate that midsagittal callosal morphology shows different trajectories for chimpanzees and humans across the adult lifespan. From young to middle adulthood, the human growth curve is steeper, reaching its maximum earlier. From middle to old adulthood, the human trajectories reveal a strong decline, that is absent in the chimpanzee sample. Thus, the over-proportionality of decline found in aging humans, is not an universal characteristic of the aging brain, and rather might represent a specific feature of the human brain. Aging studies on primates that are evolutionary more divergent from humans – like capuchin (Phillips & Sherwood, 2012) or rhesus monkeys (Bowley et al., 2010) – which also do not report significant atrophy of the corpus callosum in older age, further supporting this notion. However, the literature on age-related decline in brain measures is limited and the above-hypothesised human-specificity remains to be tested in future comparative studies.

## Supporting information

Supplementary analyses

## Acknowledgements

RW would like to thank the Institute of Psychology, University of Tartu, Estonia, for hosting his sabbatical which led to the present manuscript. The NCCC chimpanzees are supported by Cooperative Agreement U42-OD011197. The Yerkes Center and NCCC are fully accredited by the AAALAC International.

