## Supplementary analyses for "Comparative morphology of the corpus callosum across the adult lifespan in chimpanzees (Pan troglodytes) and humans"

**1. Supplement Figure S1: Age distribution**

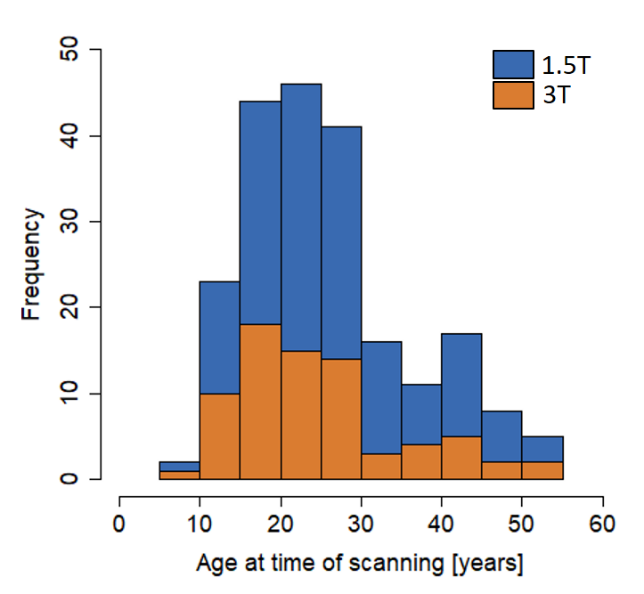

*Figure S1*. Age distribution of study sample (N=213) by MRI scanner field strength

**2. Sample selection**

The data requested from the National Chimpanzee Brain Resource (NCBR, www.chimpanzeebrain.org/) and included a total of 227 in-vivo MRI scans. Six (2.6%) were rated as “not usable” and consequently excluded from the analysis, leading to the final study sample of 221 datasets. Reasons for the low quality were high noise levels (5 cases) or corrupted data files (1 case). As indicated in the method section, also the quality of the corpus callosum segmentation was rated on a three point scale (0 = not usable, 1 = low but acceptable quality, 2 = good quality). Eight (3.5%) callosal segmentations were rated as “low quality”. Reasons were low contrast for the segmentation of the corpus callosum from fornix/blood vessels (3 cases) as well as apparent lesions within the corpus callosum (5 cases). The remaining 213 (93.8%) had “good quality” and represented the main study sample. However, for matter of completeness we repeated core analysis steps also including the “low quality” segmentation data.

**3. Human comparison data from Danielsen et al. (2020)**

The comparison samples were drawn from the sample presented in Danielsen et al. (2020) and included datasets from three imaging cohorts coordinated by the Lifespan Changes in Brain and Cognition (LCBC) research center, University of Oslo, Norway ([www.oslobrains.no](http://www.oslobrains.no)). All original cohorts excluded participants with a history of neurological conditions, any counter-indication for MRI, or non-right handedness. All original studies received ethical clearance by the Regional Ethics Committee for medical research (REK-Vest), and informed consent was given by the participants (or their guardians) for studies on brain and cognition in development and aging.

Of the total 1867 MRI datasets, 1408 (75.4% of all) were collected with a 1.5 T Avanto and 459 datasets (24.6%)using a 3 T Skyra scanners (both Siemens Medical Solutions, Erlangen, Germany) at the Oslo University Hospital, Oslo, Norway. On both systems, 3D T1-weighted magnetization prepared rapid gradient echo (MPRAGE) sequence were used. The sequence (repetition time, TR = 2400 ms; echo time, TE = 3.61 ms; inversion time, TI = 1000 ms; flip angle = 8°) used on the Avanto system covered 160 sagittal slices (1.2 mm thickness, 192 × 192 scan matrix, field of view of 240 × 240 mm^2^), resulting in a resolution of 1.25 × 1.25 × 1.20 mm^3^. The Skyra system used a comparable turbo field echo pulse sequence (TR/TE/TI = 2300 ms/2.98 ms/850 ms; flip angle = 8°) covering 176 sagittal slices (256 × 256 scan matrix) and yielding an resolution of 1.0× 1.0 × 1.0 mm^3^.

As discussed in the method section, the extraction of the initial callosal segmentation was based on white-matter segmentation using SPM12 routines (Statistical Parametric Mapping, Wellcome Department of Cognitive Neurology, London, UK) but otherwise followed the same procedures as described for the chimpanzee data. As the final segmentation was done by trained human examiners in both cases, we regard the methods leading to the segmentation as being equivalent.

**4. Comparability of chimpanzee and human age**

Human and chimpanzee lifespan development differ and in order to compare both samples directly the age range of the included human data had to be adjusted accordingly. To do so, we compared the timing of certain life events in the two species to determine a factor to transfer chimpanzee age into a human-age equivalent. The following life events were considered.

*Onset of puberty*: For male and female chimpanzees at an age of ca. 7-8 years , for example, determined based on the onset of testosterone increase (Behringer, Deschner, Deimel, Stevens, & Hohmann, 2014), while in humans an onset of ca. 10-12 years can be assumed (Behringer et al., 2014; Kelsey et al., 2014). From this a factor between 1.42 and 1.50 can be calculated.

*Age of sexual maturity*: estimated to 13.3 for chimpanzees and 19.5 for humans by Robson and Wood (Robson & Wood, 2008), suggesting a factor of 1.46.

*Maximum potential life expectancy*: chimpanzees in the wild usually die before they reach 45 years but may reach their 50s (Hill et al., 2001). In captivity, a maximum age above 60 can be observed (Edler et al., 2017). The oldest humans from hunter and forager societies live into their 70s and 80s, while an age above 120 years can be reached in modern societies (Robson & Wood, 2008). Relating the maximum life expectancy in wild chimpanzees and forger societies, as these may be considered most comparable, one yields a factor of 1.4 and 1.6.

Given the above factors range between 1.4 and 1.6, we used a factor of 1.5 in the present study to convert the age of chimpanzees to a human age equivalent. Of note, this is well in line with the factor suggested in previous comparative studies (e.g., Chen et al., 2013)

**5. R code for main GAM analyses**

The lifespan trajectories analysis of absolute and relative area and thickness analysis were fitted using the “mgcv” package (v1.8-31; (Wood, 2017) in R 3.6.2). For the main analysis the code was:

> gam.object <- gam(DV ~ s(Age, bs = "cr", k=6) + Sex + ScannerType, data = dataset, method = "REML")

Whereby DV was the respective dependent measures (e.g., relative area or relative segmental thickness).

To test for sex differences of the developmental trajectories a second analysis followed incorporating terms for the modulation of the Age trajectories by Sex. That is, the female age trajectory was fitted as reference, and deviation from this reference in males were tested. Or, expressed as R code:

> gam.object <- gam(DV ~ s(Age, bs = "cr") + s(Age, bs = "cr", by = as.ordered(Sex), k=6) + Sex + ScannerType, data = dataset, method = "REML")

Finally, difference between chimpanzee and human age trajectory (for relative area and relative thickness measures) was tested by fitting the human trajectory as reference curve, and testing for deviation in chimpanzees as follows:

> gam.object <- gam(DV ~ s(Age, bs="cr") + s(Age, bs = "cr", by= as.ordered(Species), k = 6) + Sex + ScannerType, data = dataset, method = "REML")

Of note, it was not necessary to include a “Species” predictor, as it is redundant due to its perfect collinearity with Scanner Type predictor.

**6. Supplement Figure S2: Sex differences in the ageing trajectory**

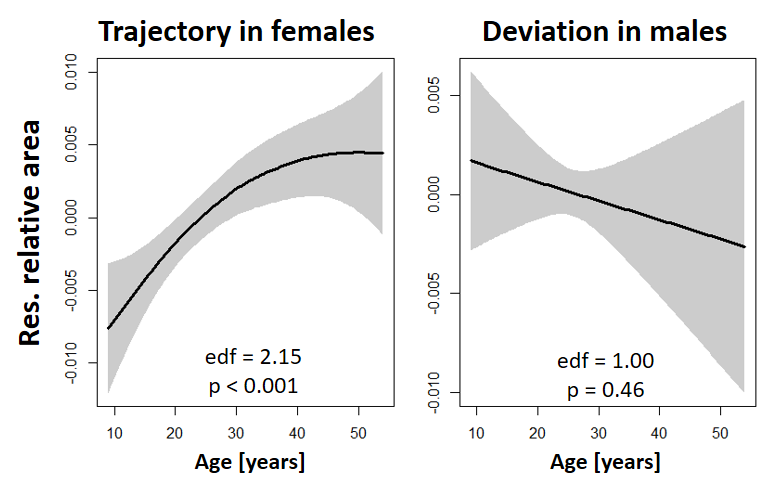

*Figure. S2.* Sex differences in aging trajectory of relative midsagittal corpus callosum. Graph shows the GAM fit (shade: 95% confidence bands) of the female trajectory (left) and the deviation from the female trajectory in males (right). While the female age trajectory was significant, the deviation in males was not significant.

**7. Supplementary analysis of absolute midsagittal area and thickness**

Considering *absolute area*, the age effect was characterised by an edf of 2.3 which was significant (*F* = 5.82, *p* < .0001) and the age predictors explained 4.9% of variance in the data (see Fig. S3 below). The slope of the trajectory was positive in young age and decreased until the age of 27.6 years where it no longer deviated from zero, marking the end of callosal growth. No decline in corpus callosum area was observed as the slope did not deviate negatively from zero in the studied age range. The main effect of Sex was not significant (*t*(207.7) = 0.77*, p* = 0.44, *ω*^2^ < .001). The follow-up analysis also did not find a deviation of the male from the female age trajectories (edf = 1.31, *F* < 1, *p* = 0.29, *ω*^2^ < 0.001).

The analysis of *absolute thickness* revealed an age effect in segments 11 to 15, with an edf of 2.32 to 2.55 and explaining between 4.1 and 6.2% of variance in the data (see Fig. S3, below). The slope of the mean trajectory within this cluster was significantly larger than zero until the age of 28 years, and the apparent negative slope in older age was not significant. For neither of the segments a main effect sex or a deviation in the trajectories between the sexes was significant (all *p_FDR_* >.30).

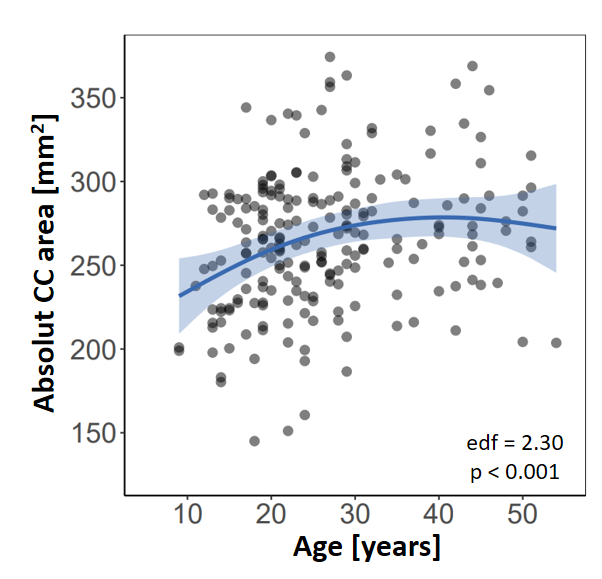

*Figure S3*. Absolute midsagittal area of the chimpanzee corpus callosum across the adult lifespan. The blue line represents the GAM fit (shading: 95% confidence band) of the prediction of relative area by Age (using Scanner type and Sex as covariate).

**8. Supplement Figure S4: Topography of edf values within the corpus callosum (and effect of inclusion of low quality data)**

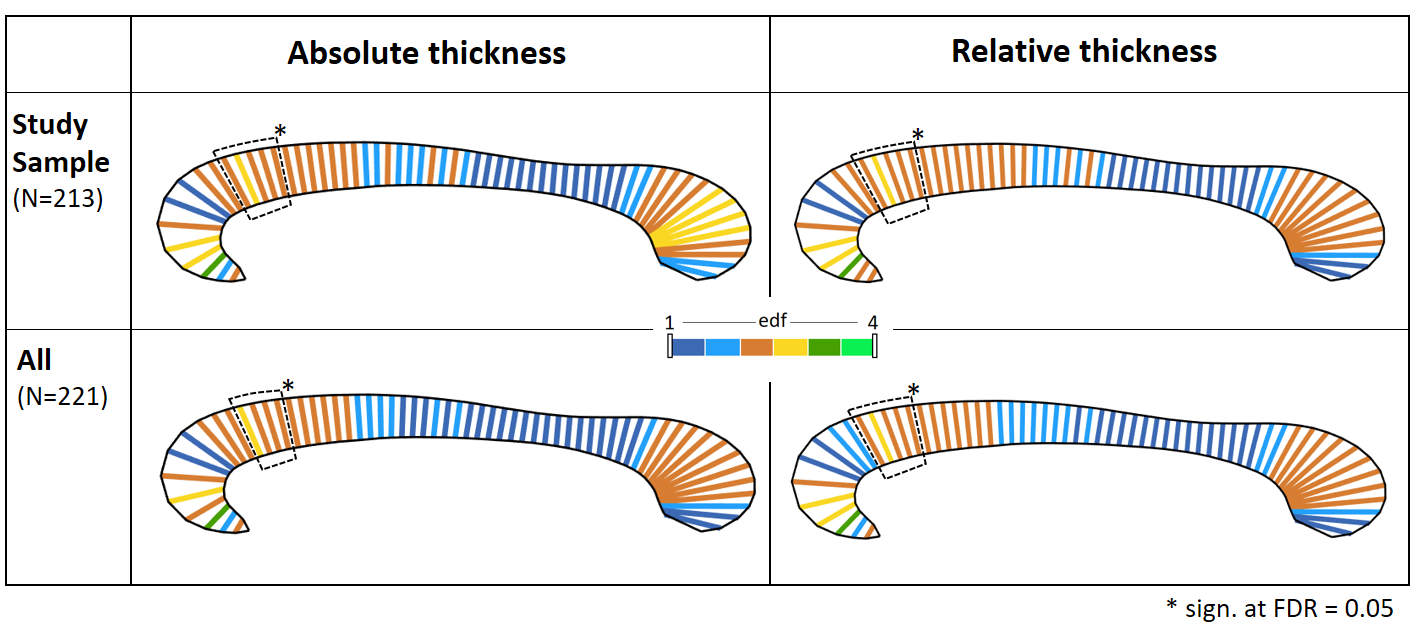

*Figure S4.* Topography of the edfs (color-coded) of the aging trajectory of relative and absolute corpus callosum thickness. Upper row shows the study sample, the lower row shows the complete sample including low quality segmentations data (for details see Supplement Section 12)

**9. Table S1. Segment-wise statistics of the relative thickness analysis**

| *Table 1* Results of the main and follow-up GAM analyses of relative thickness per segment | | | | | | | | | | |
| --- | --- | --- | --- | --- | --- | --- | --- | --- | --- | --- |
|  | Age trajectory (all)^a^ | | | Sex | Scanner |  | Age trajectory females^b^ | | Deviation in males | |
| # | *edf* | *p^c^* | *ev* | *P* | *p* |  | *edf* | *p* | *edf* | *p* |
| 01 | 2.15 | 0.33 | 0.00 | 0.94 | 0.21 |  | 2.17 | 0.41 | 1.01 | 0.91 |
| 02 | 1.77 | 0.23 | 0.01 | 0.67 | 0.97 |  | 1.00 | 0.41 | 2.52 | 0.77 |
| 03 | 3.26 | 0.15 | 0.03 | 0.34 | 0.14 |  | 3.77 | 0.31 | 1.61 | 0.77 |
| 04 | 2.48 | 0.18 | 0.02 | 0.34 | **0.03** |  | 2.13 | 0.43 | 2.04 | 0.77 |
| 05 | 2.53 | 0.07 | 0.03 | 0.54 | **0.01** |  | 2.18 | 0.35 | 2.02 | 0.77 |
| 06 | 2.04 | 0.29 | 0.00 | 0.60 | **0.01** |  | 2.02 | 0.43 | 1.00 | 0.77 |
| 07 | 1.00 | 0.18 | 0.02 | 0.34 | 0.19 |  | 1.00 | 0.37 | 1.00 | 0.77 |
| 08 | 1.00 | 0.18 | 0.02 | 0.34 | 0.64 |  | 1.00 | 0.34 | 1.00 | 0.77 |
| 09 | 2.05 | 0.21 | 0.02 | 0.38 | 0.13 |  | 2.00 | 0.35 | 1.00 | 0.77 |
| 10 | 2.03 | 0.051 | 0.04 | 0.49 | 0.15 |  | 1.99 | 0.16 | 1.00 | 0.79 |
| 11 | 2.22 | **0.02**^d^ | 0.05 | 0.74 | 0.15 |  | 2.19 | 0.11 | 1.00 | 0.78 |
| 12 | 2.50 | **0.00** | 0.06 | 0.60 | 0.21 |  | 2.52 | 0.10 | 1.00 | 0.79 |
| 13 | 2.33 | **0.00** | 0.06 | 0.67 | 0.42 |  | 2.34 | 0.10 | 1.00 | 0.77 |
| 14 | 2.32 | **0.00** | 0.07 | 0.49 | 0.71 |  | 2.34 | 0.10 | 1.00 | 0.78 |
| 15 | 2.35 | **0.00** | 0.06 | 0.83 | 0.59 |  | 2.37 | 0.10 | 1.00 | 0.87 |
| 16 | 2.24 | 0.055 | 0.03 | 0.65 | 0.79 |  | 2.26 | 0.11 | 1.00 | 0.87 |
| 17 | 2.36 | 0.06 | 0.03 | 0.34 | 0.91 |  | 2.41 | 0.11 | 1.00 | 0.97 |
| 18 | 2.23 | 0.20 | 0.02 | 0.37 | 0.70 |  | 2.25 | 0.25 | 1.00 | 0.87 |
| 19 | 2.06 | 0.24 | 0.01 | 0.35 | 0.22 |  | 2.07 | 0.39 | 1.00 | 0.78 |
| 20 | 2.21 | 0.22 | 0.01 | 0.24 | 0.20 |  | 2.24 | 0.29 | 1.00 | 0.87 |
| 21 | 2.20 | 0.24 | 0.01 | 0.17 | 0.18 |  | 2.23 | 0.29 | 1.00 | 0.97 |
| 22 | 1.90 | 0.29 | 0.00 | 0.17 | 0.30 |  | 1.90 | 0.41 | 1.00 | 0.87 |
| 23 | 1.61 | 0.53 | 0.00 | 0.08 | 0.12 |  | 1.63 | 0.54 | 1.00 | 0.87 |
| 24 | 1.74 | 0.45 | 0.00 | 0.05 | 0.05 |  | 1.75 | 0.48 | 1.00 | 0.87 |
| 25 | 1.79 | 0.44 | 0.00 | 0.09 | **0.01** |  | 1.82 | 0.42 | 1.00 | 0.87 |
| 26 | 1.47 | 0.47 | 0.00 | 0.24 | **0.01** |  | 1.49 | 0.48 | 1.00 | 0.87 |
| 27 | 1.45 | 0.47 | 0.00 | 0.25 | **0.01** |  | 1.30 | 0.48 | 1.29 | 0.87 |
| 28 | 1.38 | 0.45 | 0.00 | 0.15 | **0.00** |  | 1.00 | 0.41 | 1.86 | 0.78 |
| 29 | 1.85 | 0.33 | 0.00 | 0.15 | **0.01** |  | 1.41 | 0.41 | 1.85 | 0.77 |
| 30 | 1.27 | 0.55 | 0.00 | 0.17 | **0.00** |  | 1.00 | 0.41 | 2.08 | 0.77 |
| 31 | 1.79 | 0.45 | 0.00 | 0.16 | **0.01** |  | 1.00 | 0.48 | 2.24 | 0.77 |
| 32 | 1.19 | 0.61 | 0.00 | 0.17 | **0.01** |  | 1.00 | 0.59 | 1.89 | 0.78 |
| 33 | 1.00 | 0.85 | 0.00 | **0.048** | **0.01** |  | 1.00 | 0.85 | 1.75 | 0.79 |
| 34 | 1.00 | 0.45 | 0.00 | **0.048** | **0.01** |  | 1.00 | 0.63 | 1.79 | 0.79 |
| 35 | 1.00 | 0.45 | 0.00 | **0.048** | **0.04** |  | 1.00 | 0.70 | 1.00 | 0.79 |
| 36 | 1.00 | 0.45 | 0.00 | 0.09 | **0.03** |  | 1.00 | 0.60 | 1.00 | 0.87 |
| 37 | 1.00 | 0.23 | 0.01 | 0.17 | 0.06 |  | 1.00 | 0.19 | 1.00 | 0.77 |
| 38 | 1.00 | 0.20 | 0.01 | 0.17 | 0.27 |  | 1.00 | 0.12 | 1.00 | 0.77 |
| 39 | 1.00 | 0.22 | 0.01 | 0.16 | 0.31 |  | 1.00 | 0.11 | 1.46 | 0.77 |
| 40 | 1.00 | 0.24 | 0.01 | 0.24 | 0.63 |  | 1.00 | 0.11 | 1.25 | 0.77 |
| 41 | 1.00 | 0.25 | 0.01 | 0.49 | 0.97 |  | 1.00 | 0.11 | 1.00 | 0.77 |
| 42 | 1.13 | 0.30 | 0.00 | 0.41 | 0.78 |  | 1.00 | 0.11 | 1.00 | 0.43 |
| 43 | 1.00 | 0.29 | 0.00 | 0.41 | 0.68 |  | 1.00 | 0.11 | 1.00 | 0.59 |
| 44 | 1.00 | 0.24 | 0.01 | 0.74 | 0.91 |  | 1.00 | 0.15 | 1.00 | 0.77 |
| 45 | 1.00 | 0.24 | 0.01 | 0.95 | 0.72 |  | 1.00 | 0.16 | 1.00 | 0.77 |
| 46 | 1.00 | 0.25 | 0.01 | 0.73 | 0.76 |  | 1.00 | 0.11 | 1.12 | 0.77 |
| 47 | 1.39 | 0.29 | 0.01 | 0.49 | 0.60 |  | 1.61 | 0.20 | 1.04 | 0.77 |
| 48 | 1.82 | 0.33 | 0.00 | 0.98 | 0.72 |  | 2.02 | 0.30 | 1.00 | 0.77 |
| 49 | 2.31 | 0.23 | 0.02 | 0.65 | 0.22 |  | 2.52 | 0.22 | 1.00 | 0.79 |
| 50 | 2.21 | 0.26 | 0.01 | 0.49 | 0.19 |  | 2.31 | 0.29 | 1.00 | 0.87 |
| 51 | 2.43 | 0.23 | 0.01 | 0.34 | 0.07 |  | 2.66 | 0.22 | 1.00 | 0.77 |
| 52 | 2.41 | 0.24 | 0.01 | 0.25 | **0.02** |  | 2.60 | 0.25 | 1.00 | 0.78 |
| 53 | 2.49 | 0.23 | 0.01 | 0.22 | **0.02** |  | 2.58 | 0.24 | 1.03 | 0.87 |
| 54 | 2.46 | 0.22 | 0.01 | 0.17 | **0.01** |  | 2.54 | 0.22 | 1.00 | 0.79 |
| 55 | 2.42 | 0.22 | 0.02 | 0.42 | **0.01** |  | 2.48 | 0.25 | 1.27 | 0.77 |
| 56 | 2.37 | 0.23 | 0.01 | 0.95 | **0.00** |  | 2.36 | 0.29 | 1.59 | 0.77 |
| 57 | 2.21 | 0.28 | 0.01 | 0.41 | **0.00** |  | 2.31 | 0.29 | 1.00 | 0.79 |
| 58 | 1.73 | 0.56 | 0.00 | 0.27 | **0.00** |  | 1.77 | 0.59 | 1.00 | 0.90 |
| 59 | 1.01 | 0.94 | 0.00 | 0.05 | **0.01** |  | 1.00 | 0.83 | 1.00 | 0.87 |
| 60 | 1.06 | 0.35 | 0.00 | 0.24 | **0.01** |  | 1.20 | 0.29 | 1.00 | 0.77 |
| *Notes.* (a) main analysis; (b) follow-up analysis: differential sex effects; (c) all p values are corrected to achieve FDR = 0.05; (d) significant effect are highlighted with bold font | | | | | | | | | | |

**10. Table S2: Segment-wise statistics for comparison of chimpanzee with human aging trajectories of corpus callosum thickness**

| *Table S3*. Results of segment-by-segment comparison between human and chimpanzee age trajectories in relative thickness for chronological-age and adjusted-age comparisons | | | | | | | | | | | | | |
| --- | --- | --- | --- | --- | --- | --- | --- | --- | --- | --- | --- | --- | --- |
|  | Chronological-age comparison | | | | | |  | Adjusted-age comparison | | | | | |
|  | Trajectory humans | | | Deviation chimpanzees | | |  | Trajectory humans | | | Deviation chimpanzees | | |
| # | *edf* | *p^a^* | *ev^b^* | *edf* | *p* | *ev* |  | *edf* | *p* | *ev* | *edf* | *p* | *ev* |
| 01 | 1.00 | **<0.001** | 0.03 | 1.89 | **0.03** | 0.01 |  | 1.58 | **<0.001** | 0.04 | 2.11 | **0.002** | 0.01 |
| 02 | 1.00 | **<0.001** | 0.05 | 1.59 | **<0.001** | 0.02 |  | 2.51 | **<0.001** | 0.06 | 2.26 | **<0.001** | 0.02 |
| 03 | 1.00 | **<0.001** | 0.04 | 3.59 | **<0.001** | 0.03 |  | 2.88 | **<0.001** | 0.04 | 3.38 | **<0.001** | 0.03 |
| 04 | 1.00 | **<0.001** | 0.04 | 2.79 | **<0.001** | 0.03 |  | 2.79 | **<0.001** | 0.04 | 2.98 | **<0.001** | 0.03 |
| 05 | 1.00 | **<0.001** | 0.03 | 2.63 | **<0.001** | 0.03 |  | 2.42 | **<0.001** | 0.03 | 2.81 | **<0.001** | 0.03 |
| 06 | 1.00 | **0.01** | 0.01 | 2.00 | 0.06 | 0.01 |  | 1.01 | **<0.001** | 0.05 | 1.99 | **<0.001** | 0.01 |
| 07 | 1.00 | 0.61 | 0.00 | 1.00 | 0.23 | 0.00 |  | 3.29 | **<0.001** | 0.04 | 1.86 | **<0.001** | 0.02 |
| 08 | 1.00 | 0.14 | 0.00 | 1.00 | 0.43 | 0.00 |  | 3.56 | **<0.001** | 0.04 | 2.05 | **<0.001** | 0.02 |
| 09 | 2.25 | 0.32 | 0.00 | 1.06 | 0.34 | 0.00 |  | 3.42 | **<0.001** | 0.04 | 1.00 | **<0.001** | 0.03 |
| 10 | 1.75 | 0.61 | 0.00 | 1.57 | 0.09 | 0.01 |  | 3.50 | **<0.001** | 0.04 | 1.01 | **<0.001** | 0.04 |
| 11 | 1.96 | 0.55 | 0.00 | 1.77 | **0.03** | 0.01 |  | 3.52 | **<0.001** | 0.05 | 1.00 | **<0.001** | 0.04 |
| 12 | 1.68 | 0.70 | 0.00 | 2.36 | **0.01** | 0.01 |  | 3.37 | **<0.001** | 0.05 | 1.00 | **<0.001** | 0.05 |
| 13 | 1.01 | 0.60 | 0.00 | 2.42 | **0.00** | 0.02 |  | 3.37 | **<0.001** | 0.05 | 1.00 | **<0.001** | 0.05 |
| 14 | 1.00 | 0.73 | 0.00 | 2.42 | **0.00** | 0.02 |  | 3.45 | **<0.001** | 0.05 | 1.00 | **<0.001** | 0.06 |
| 15 | 1.00 | 0.43 | 0.00 | 2.45 | **0.00** | 0.02 |  | 3.67 | **<0.001** | 0.05 | 1.00 | **<0.001** | 0.05 |
| 16 | 1.00 | 0.29 | 0.00 | 2.36 | **0.00** | 0.02 |  | 3.73 | **<0.001** | 0.06 | 1.00 | **<0.001** | 0.04 |
| 17 | 1.00 | 0.32 | 0.00 | 2.45 | **0.00** | 0.02 |  | 3.55 | **<0.001** | 0.05 | 1.01 | **<0.001** | 0.04 |
| 18 | 1.00 | 0.43 | 0.00 | 2.33 | **0.03** | 0.01 |  | 3.28 | **<0.001** | 0.04 | 1.00 | **<0.001** | 0.03 |
| 19 | 1.00 | 0.55 | 0.00 | 2.25 | 0.06 | 0.01 |  | 3.42 | **<0.001** | 0.05 | 1.00 | **<0.001** | 0.03 |
| 20 | 1.00 | 0.43 | 0.00 | 2.40 | 0.03^c^ | 0.01 |  | 3.19 | **<0.001** | 0.04 | 1.02 | **<0.001** | 0.03 |
| 21 | 1.00 | 0.76 | 0.00 | 2.33 | 0.08 | 0.01 |  | 3.36 | **<0.001** | 0.04 | 1.00 | **<0.001** | 0.02 |
| 22 | 1.00 | 0.75 | 0.00 | 2.04 | 0.13 | 0.00 |  | 3.19 | **<0.001** | 0.05 | 1.00 | **<0.001** | 0.03 |
| 23 | 1.00 | 0.97 | 0.00 | 1.75 | 0.40 | 0.00 |  | 2.84 | **<0.001** | 0.04 | 1.00 | **<0.001** | 0.02 |
| 24 | 1.03 | 0.55 | 0.00 | 1.80 | 0.41 | 0.00 |  | 3.14 | **<0.001** | 0.04 | 1.00 | **<0.001** | 0.02 |
| 25 | 1.00 | 0.73 | 0.00 | 1.93 | 0.32 | 0.00 |  | 3.00 | **<0.001** | 0.03 | 1.00 | **<0.001** | 0.02 |
| 26 | 1.93 | 0.55 | 0.00 | 1.09 | 0.42 | 0.00 |  | 2.77 | **<0.001** | 0.03 | 1.00 | **<0.001** | 0.02 |
| 27 | 1.72 | 0.65 | 0.00 | 1.34 | 0.53 | 0.00 |  | 2.94 | **<0.001** | 0.02 | 1.00 | **<0.001** | 0.01 |
| 28 | 2.06 | 0.43 | 0.00 | 1.01 | 0.48 | 0.00 |  | 2.93 | **<0.001** | 0.02 | 1.00 | **<0.001** | 0.02 |
| 29 | 1.82 | 0.43 | 0.00 | 1.64 | 0.47 | 0.00 |  | 3.04 | **<0.001** | 0.02 | 1.00 | **<0.001** | 0.02 |
| 30 | 2.06 | 0.32 | 0.00 | 1.00 | 0.72 | 0.00 |  | 2.89 | **<0.001** | 0.02 | 1.00 | **0.001** | 0.01 |
| 31 | 1.89 | 0.32 | 0.00 | 1.50 | 0.71 | 0.00 |  | 3.17 | **<0.001** | 0.02 | 1.00 | **0.002** | 0.01 |
| 32 | 1.25 | **0.04** | 0.01 | 1.15 | 0.83 | 0.00 |  | 3.15 | **0.001** | 0.01 | 1.00 | **0.02** | 0.01 |
| 33 | 1.01 | **0.02** | 0.01 | 1.00 | 0.53 | 0.00 |  | 3.10 | **0.01** | 0.01 | 1.01 | 0.08 | 0.00 |
| 34 | 1.14 | **0.01** | 0.01 | 1.00 | 0.77 | 0.00 |  | 3.16 | **0.01** | 0.01 | 1.16 | **0.04** | 0.00 |
| 35 | 1.35 | **0.01** | 0.01 | 1.00 | 0.75 | 0.00 |  | 3.23 | **0.002** | 0.01 | 1.67 | **0.04** | 0.00 |
| 36 | 2.42 | 0.10 | 0.01 | 1.00 | 0.99 | 0.00 |  | 2.84 | **0.003** | 0.01 | 1.27 | **0.03** | 0.00 |
| 37 | 2.17 | 0.32 | 0.00 | 1.00 | 0.40 | 0.00 |  | 2.51 | **0.004** | 0.01 | 1.00 | **0.001** | 0.01 |
| 38 | 1.42 | 0.73 | 0.00 | 1.00 | 0.14 | 0.00 |  | 2.23 | **0.005** | 0.01 | 1.01 | **0.001** | 0.01 |
| 39 | 1.00 | 0.54 | 0.00 | 1.00 | 0.06 | 0.01 |  | 2.33 | **<0.001** | 0.02 | 1.00 | **<0.001** | 0.02 |
| 40 | 1.00 | 0.15 | 0.00 | 1.00 | **0.03** | 0.01 |  | 1.20 | **<0.001** | 0.02 | 1.00 | **<0.001** | 0.02 |
| 41 | 1.00 | **0.05** | 0.01 | 1.00 | **0.03** | 0.01 |  | 1.01 | **<0.001** | 0.04 | 1.01 | **<0.001** | 0.02 |
| 42 | 1.42 | **0.04** | 0.01 | 1.04 | **0.02** | 0.01 |  | 1.01 | **<0.001** | 0.04 | 1.33 | **<0.001** | 0.01 |
| 43 | 1.04 | **0.01** | 0.01 | 1.00 | **0.02** | 0.01 |  | 2.61 | **<0.001** | 0.05 | 1.00 | **<0.001** | 0.02 |
| 44 | 1.00 | **0.01** | 0.01 | 1.00 | **0.02** | 0.01 |  | 3.13 | **<0.001** | 0.05 | 1.00 | **<0.001** | 0.02 |
| 45 | 1.00 | **0.01** | 0.01 | 1.00 | **0.02** | 0.01 |  | 2.92 | **<0.001** | 0.05 | 1.00 | **<0.001** | 0.02 |
| 46 | 1.00 | **0.03** | 0.01 | 1.00 | **0.03** | 0.01 |  | 2.75 | **<0.001** | 0.05 | 1.00 | **<0.001** | 0.02 |
| 47 | 1.00 | 0.10 | 0.00 | 1.54 | 0.10 | 0.01 |  | 2.88 | **<0.001** | 0.05 | 1.00 | **<0.001** | 0.02 |
| 48 | 1.13 | 0.29 | 0.00 | 1.93 | 0.16 | 0.00 |  | 3.02 | **<0.001** | 0.05 | 1.00 | **<0.001** | 0.02 |
| 49 | 1.81 | 0.54 | 0.00 | 2.28 | 0.20 | 0.00 |  | 3.06 | **<0.001** | 0.05 | 1.00 | **<0.001** | 0.03 |
| 50 | 2.25 | 0.43 | 0.00 | 2.03 | 0.37 | 0.00 |  | 3.19 | **<0.001** | 0.06 | 1.00 | **<0.001** | 0.02 |
| 51 | 2.32 | 0.43 | 0.00 | 2.31 | 0.40 | 0.00 |  | 3.29 | **<0.001** | 0.06 | 1.00 | **<0.001** | 0.02 |
| 52 | 2.74 | 0.26 | 0.00 | 2.16 | 0.54 | 0.00 |  | 3.43 | **<0.001** | 0.06 | 1.00 | **<0.001** | 0.02 |
| 53 | 2.97 | 0.18 | 0.00 | 2.15 | 0.43 | 0.00 |  | 3.55 | **<0.001** | 0.06 | 1.00 | **0.001** | 0.01 |
| 54 | 3.24 | 0.08 | 0.01 | 1.97 | 0.43 | 0.00 |  | 3.50 | **<0.001** | 0.05 | 1.00 | **0.002** | 0.01 |
| 55 | 3.45 | **0.01** | 0.01 | 1.80 | 0.27 | 0.00 |  | 3.53 | **<0.001** | 0.04 | 1.00 | **0.03** | 0.00 |
| 56 | 3.44 | **0.00** | 0.02 | 1.49 | 0.31 | 0.00 |  | 3.82 | **<0.001** | 0.03 | 1.00 | 0.06 | 0.00 |
| 57 | 3.72 | **0.00** | 0.05 | 1.00 | 0.09 | 0.00 |  | 4.06 | **<0.001** | 0.03 | 1.00 | 0.45 | 0.00 |
| 58 | 3.75 | **0.00** | 0.06 | 1.00 | 0.01^c^ | 0.01 |  | 3.12 | **<0.001** | 0.03 | 1.00 | 0.09 | 0.00 |
| 59 | 3.95 | **0.00** | 0.06 | 1.00 | 0.01^c^ | 0.01 |  | 2.32 | **<0.001** | 0.05 | 1.00 | **0.001** | 0.01 |
| 60 | 2.68 | **0.00** | 0.04 | 1.00 | 0.06 | 0.00 |  | 1.48 | **<0.001** | 0.05 | 1.00 | **0.02** | 0.01 |
| *Notes.* (a) all *p* values are corrected to achieve FDR = 0.05; (b) explained variance effect; (c) segment significant, but excluded due to extent threshold (see text for details) | | | | | | | | | | | | | |

**11. Supplement Figure S5: Topography of the age trajectories in human samples**

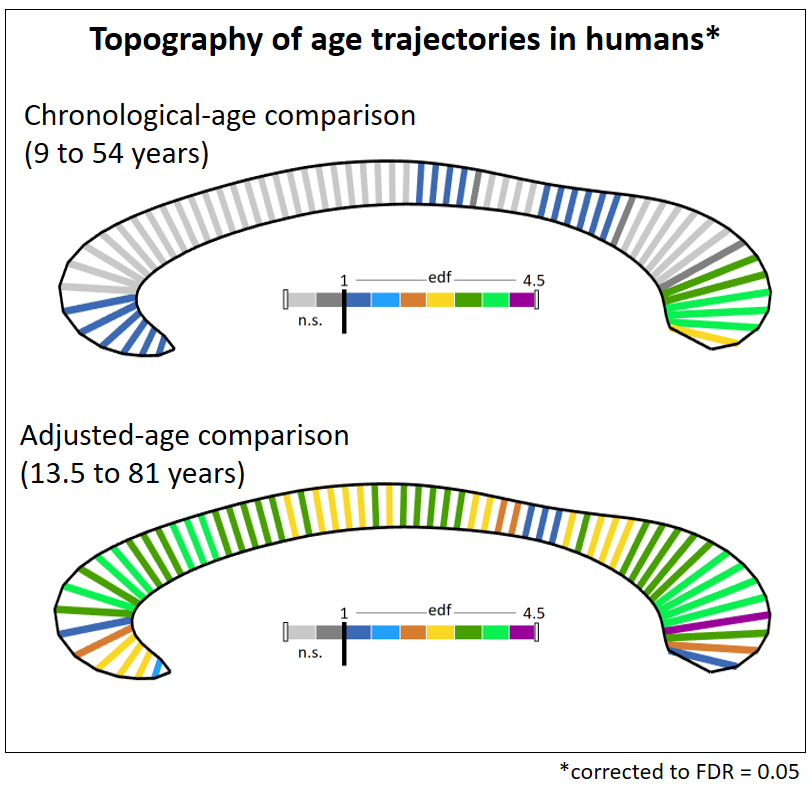

*Figure S5.* Topography of the edfs (color-coded) of the aging trajectory of relative corpus callosum thickness. The upper panel represents the trajectory in the chronological-age reference sample, and the lower panel the trajectories in the adjusted-age sample. The effects are stronger in the adjusted-age sample, as here the human sample includes participants above the age of 60 for which the atrophy of the corpus callosum is known to be strongest (Danielsen et al., 2020)

**12. Analysis of age trajectories including all low quality data sets (N=221).**

Adding the eight participants of “low quality” callosal segmentation does not substantially change the results compared to when these datasets were excluded. Including the 8 additional subjects, yields a sample 221 data sets (135 from females) with a mean age of 27.0 ± 10.4 years (covering an age range from 9 to 54 years). The results for this subsample were as follows:

*Absolute corpus callosum area*: The fitted age trajectory had an edf of 2.3 which was significant (*p* = .002, *ω*^2^ < 0.042). The main effect of Sex was not significant (*t*(215.7) = 0.92*, p* = 0.36, *ω*^2^ < 0.001). A direct comparison of the trajectories did not find a deviation in males from females (edf = 1.17, *p* = 0.34, *ω*^2^ < 0.001).

*Relative corpus callosum area*: The age trajectory had an edf of 2.11 which was significant (*p* < .001, *ω*^2^ < 0.066). The main effect of Sex was not significant (*t*(215.9) = -1.15*, p* = 0.25, *ω*^2^ < 0.01) and the trajectories did not differ between the sexes (deviation: edf = 1.00, *p* = 0.58, *ω*^2^ < 0.01).

*Callosal thickness analysis*: As can be seen in Fig. S4, the topography of the edf values and the regions that show significant age effects are comparable for both samples. The only obvious difference was that regarding absolute thickness measures the trajectory of segment 12 did not survive FDR correction in the full sample, while it did in the study sample.

**13. Trajectory analysis in human sample**

In order to determine end of growth and beginning of decline of relative corpus callosum area in humans, we ran a GAM analysis of the human sample analogue to the analysis conducted in chimpanzees (see R code in Supplement Section 5). For comparability, we restricted the human sample to all first-time scanning data to yield a purely cross-sectional analysis. The resulting sample consisted of 1016 datasets (age range: 4.1 to 93 years; n = 609 or 59.9% were female). The age fit was significant with an edf = 4.15 (*F* = 56.0, *p* < .0001; *ω*^2^ = .18), indicating an inverted u-shape association of age and relative callosal thickness (see Figure S6, below). We then determined the derivatives (i.e., slope) of the fitted trajectories to estimate the age at which the slope is the last time above and the first time below zero (using the 95% confidence bands), defining the end of growth and the beginning of decline, respectively. That is, the end of growth was estimated to be at an age of 23 years and the beginning of decline was found to be at an age of 51.0 years.

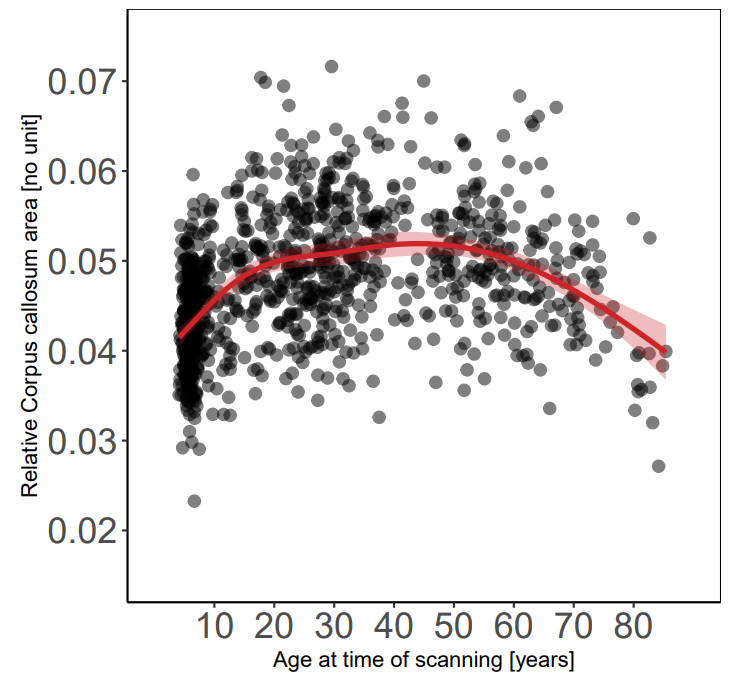

*Figure S6*. Fitted relative midsagittal area of the human corpus callosum across the lifespan. The red line represents the GAM fit (shading: 95% confidence band) of the prediction of relative area by Age (using Scanner type and Sex as covariates).
